## Supplementary figures and images for "Time-resolved cryo-EM visualizes ribosomal translocation with EF-G and GTP"

### Supplementary Movie 1

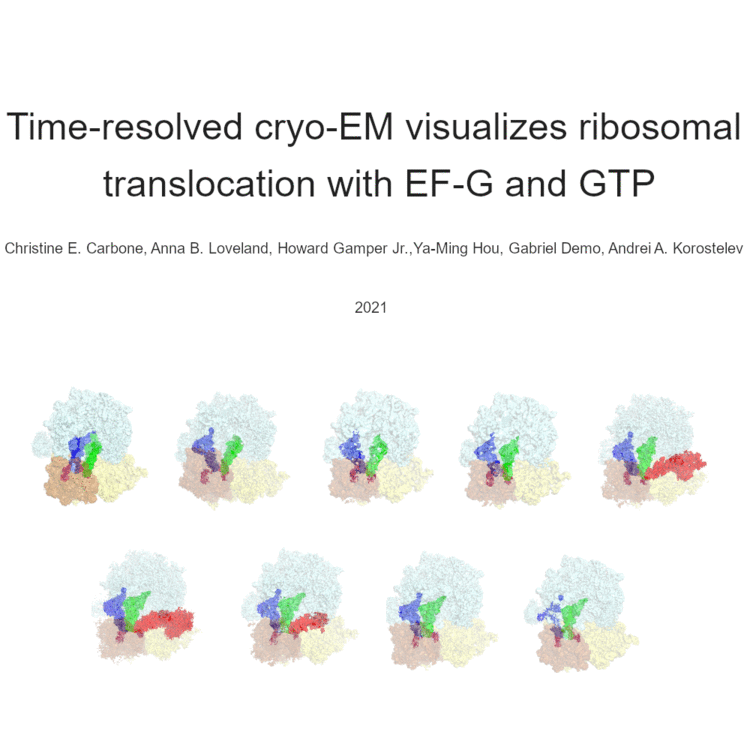
